## Supplementary material for "H/ACA snoRNP guides ribosomal RNA subdomain folding in a satellite particle before joining the core 90S pre-ribosome": Document S1

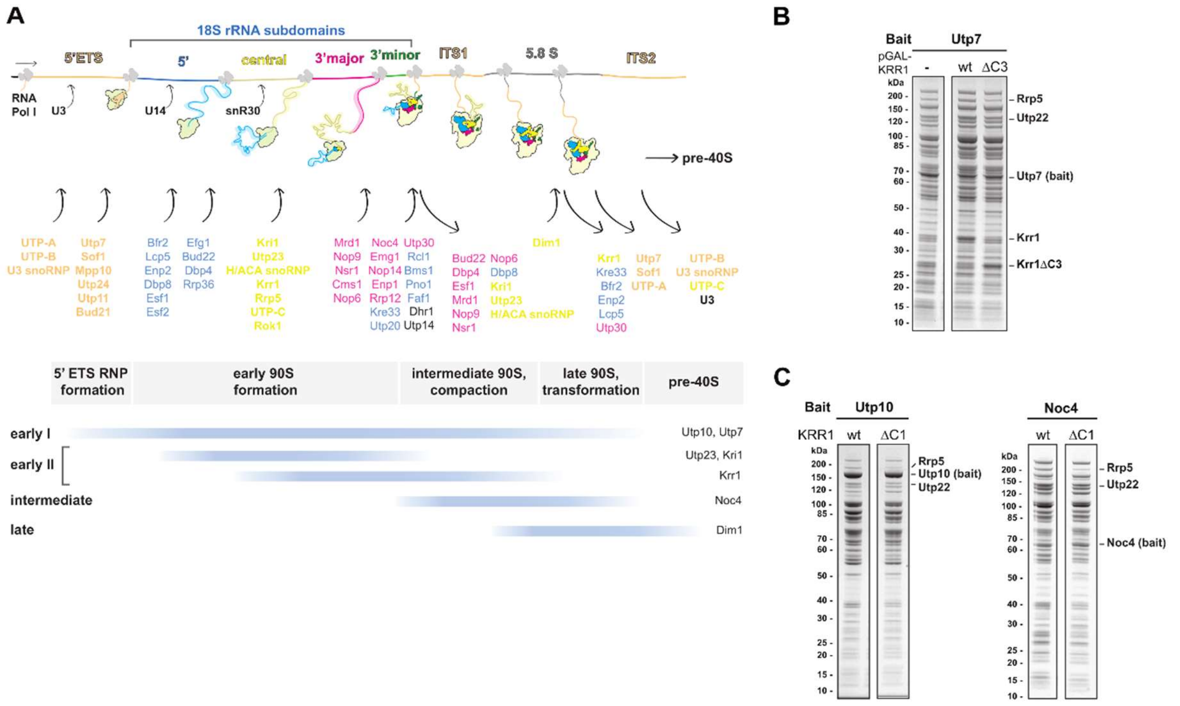


**Figure S1 (related to Figure 1). Formation and maturation of 90S pre-ribosomes.** (**A**) Schematic overview of 90S pre-ribosome biogenesis. AFs are recruited to the evolving 90S pre-ribosome co-transcriptionally to specific 18S pre-rRNA subdomains. Timeline is based on ([9-11]) and ([5,7,8]). (**B**) SDS-PAGE of early 90S particles isolated via Utp7 from cells grown in galactose containing medium either harboring an empty plasmid (-) or glucose inducible wild-type *KRR1* or *krr1*ΔC3. (**C**) SDS-PAGE of early (Utp10) and intermediate (Noc4) 90S particles isolated from wild-type or *krr1*ΔC1 cells.

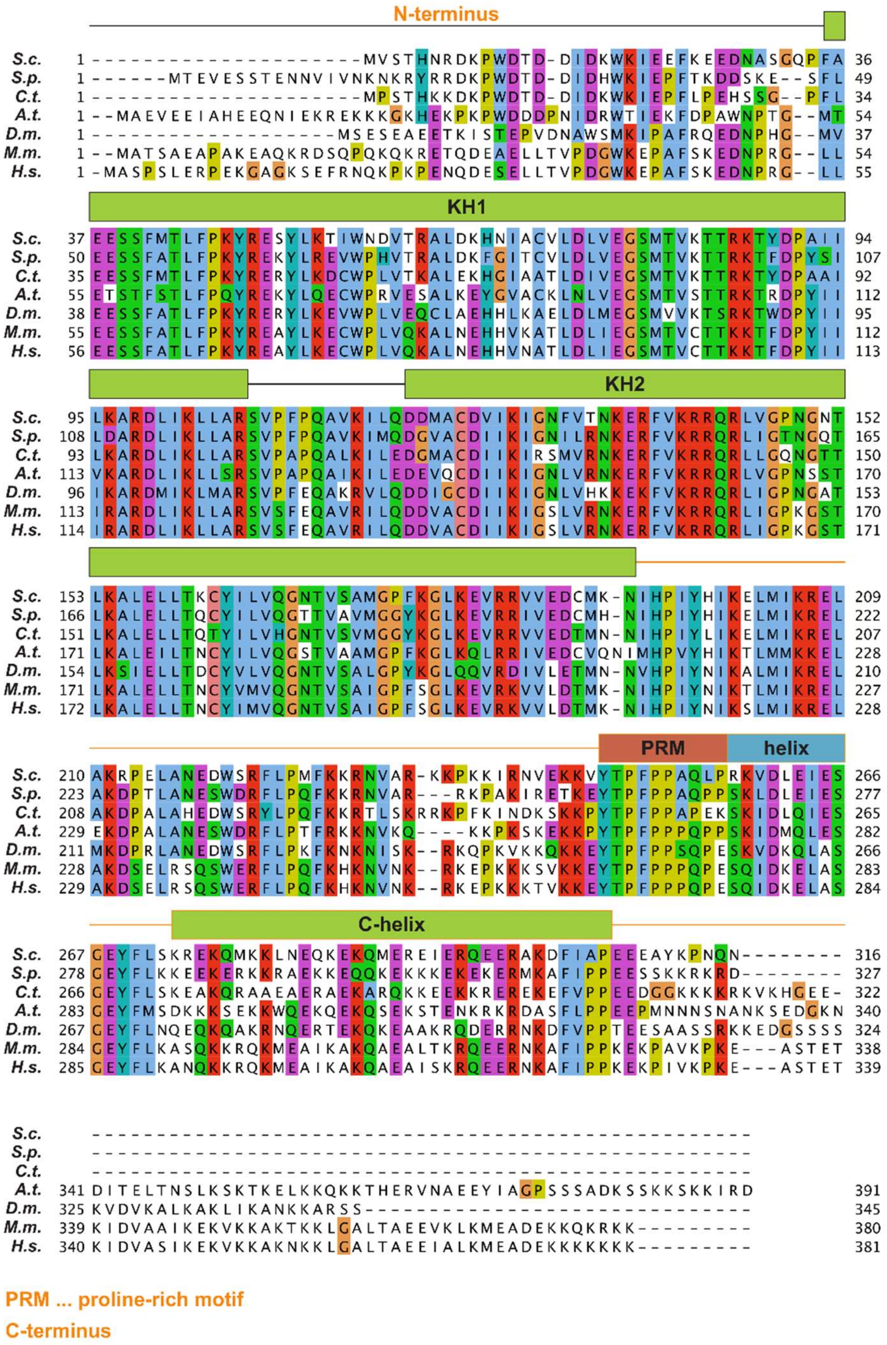


**Figure S2 (related to Figure 1). Krr1 harbors a conserved proline-rich motif within its C-terminal domain.** Multiple sequence alignment of Krr1. C-terminal truncation mutants used in this study as well as the long C-helix and the proline-rich motif are indicated. Sequences of *S.c. Saccharomyces cerevisiae*, *S.p. Schizosaccharomyces pombe*, *C.t. Chaetomium thermophilum*, *A.t. Arabidopsis thaliana*, *D.m. Drosophila melanogaster*, *M.m. Mus musculus*, *H.s. Homo sapiens* were aligned with Clustal Omega and displayed with Jalview.

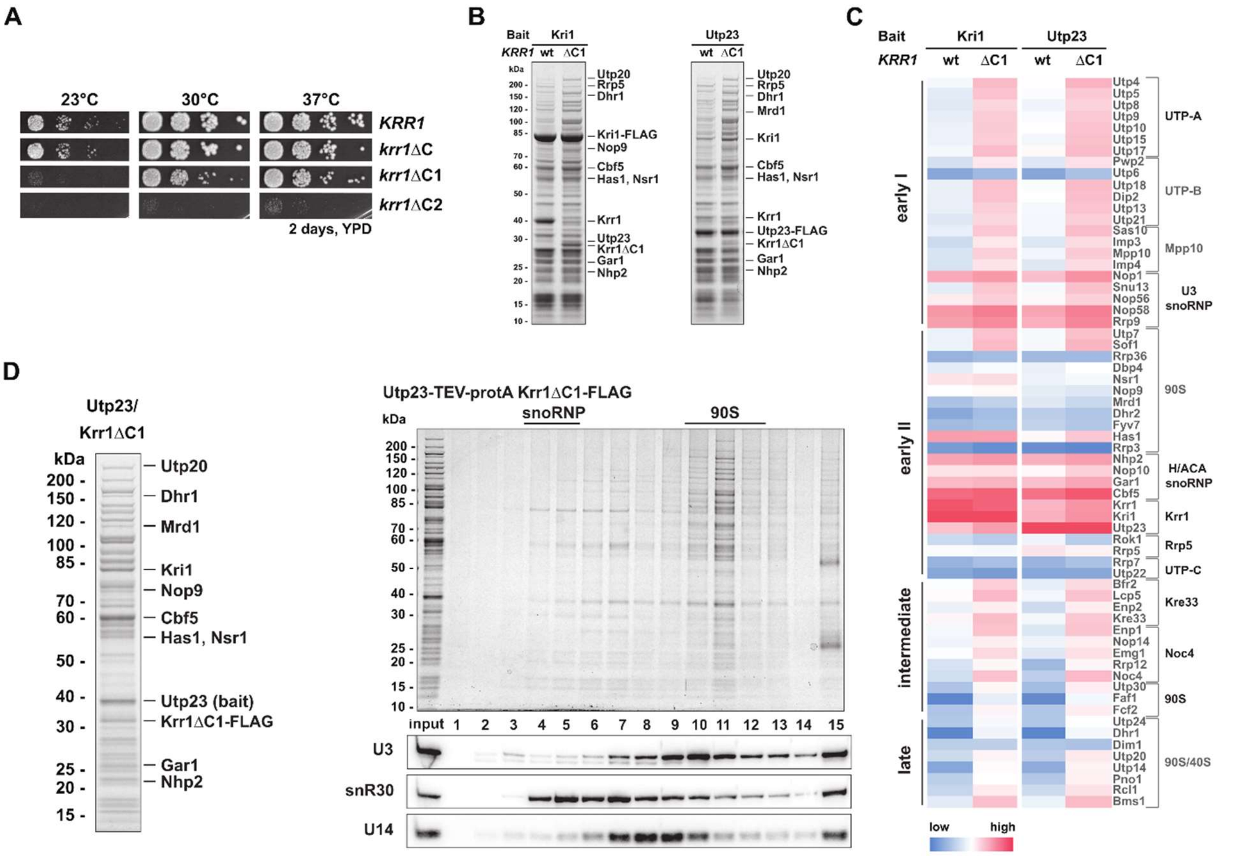


**Figure S3 (related to Figure 2). Conserved proline-rich motif of Krr1 affects association of Utp23 and Kri1 with 90S particles.** (**A**) Growth analysis of viable Krr1-CTD truncation mutants at different temperatures. (**B**) SDS-PAGE of proteins associated with Kri1 and Utp23 from wt and *krr1*ΔC1 mutant cells. (**C**) Heat map of the proteins associated with Kri1 and Utp23 from wt and *krr1*ΔC1 cells as shown in (B). Intensity-based iBAQ values were normalized to the bait, and $\log_{10}$ values of the normalized iBAQ values were colored from low (blue) to high (red). Identified proteins are grouped according to their 90S biogenesis module on the right, or according to their dynamic association (Figure S1A) on the left. (**D**) SDS-PAGE of proteins isolated via Utp23-Krr1ΔC1 (**E**) Sucrose gradient analysis of the Utp23-Krr1ΔC1 purification and northern blot analysis of sucrose gradient fractions with probes targeting snoRNAs snR30, U3 and U14.

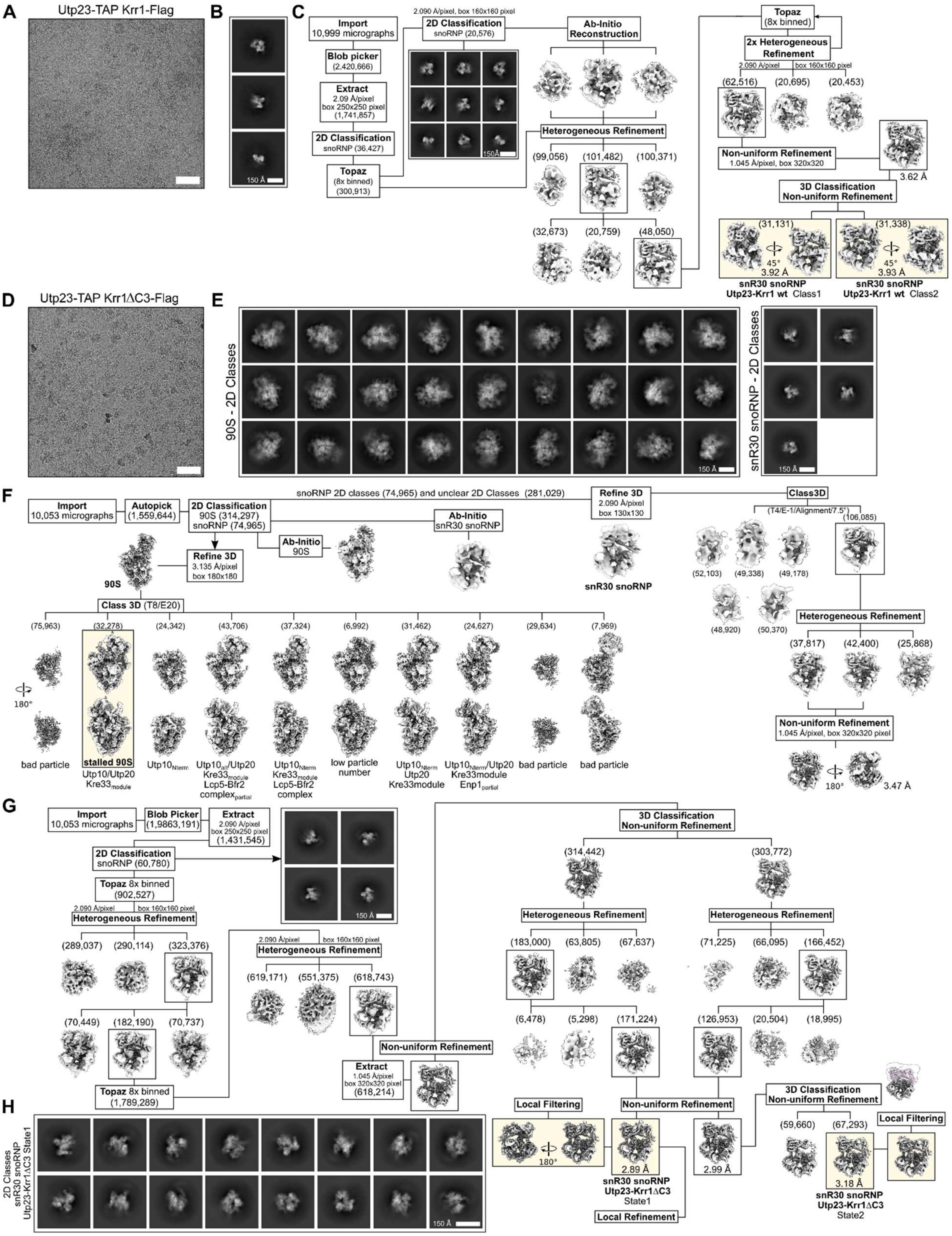
A
Utp23-TAP Krr1-Flag
B
150 Å
C
Import
10,999 micrographs
Blob picker
(2,420,666)
Extract
2.09 Å/pixel
box 250x250 pixel
(1,741,857)
2D Classification
snoRNP (36,427)
Topaz
(8x binned)
(300,913)
2.090 Å/pixel, box 160x160 pixel
2D Classification
snoRNP (20,576)
Ab-Initio
Reconstruction
Heterogeneous Refinement
(99,056)
(101,482)
(100,371)
(32,673)
(20,759)
(48,050)
Topaz
(8x binned)
2x Heterogeneous
Refinement
2.090 Å/pixel
box 160x160 pixel
(62,516)
(20,695)
(20,453)
Non-uniform Refinement
1.045 Å/pixel, box 320x320
3.62 Å
3D Classification
Non-uniform Refinement
(31,131)
(31,338)
45°
3.92 Å
3.93 Å
snR30 snoRNP
Utp23-Krr1 wt Class1
snR30 snoRNP
Utp23-Krr1 wt Class2
D
Utp23-TAP Krr1ΔC3-Flag
E
90S - 2D Classes
snR30 snoRNP - 2D Classes
150 Å
F
Import
10,053 micrographs
Autopick
(1,559,644)
2D Classification
90S (314,297)
snoRNP (74,965)
snoRNP 2D classes (74,965) and unclear 2D Classes (281,029)
Refine 3D
2.090 Å/pixel
box 130x130
Class3D
(T4/E-1/Alignment/7.5°)
Ab-Initio
snR30 snoRNP
Ab-Initio
90S
Refine 3D
3.135 Å/pixel
box 180x180
90S
snR30 snoRNP
(52,103)
(49,338)
(49,178)
(106,085)
(48,920)
(50,370)
Class 3D (T8/E20)
(75,963)
(32,278)
(24,342)
(43,706)
(37,324)
(6,992)
(31,462)
(24,627)
(29,634)
(7,969)
180°
bad particle
stalled 90S
Utp10/Utp20
Kre33module
Utp10Nterm
Utp10all/Utp20
Kre33module
Lcp5-Bfr2
complexpartial
Utp10Nterm
Kre33module
Lcp5-Bfr2
complex
low particle
number
Utp10Nterm
Utp20
Kre33module
Utp10Nterm/Utp20
Kre33module
Enp1partial
bad particle
bad particle
Heterogeneous Refinement
(37,817)
(42,400)
(25,868)
Non-uniform Refinement
1.045 Å/pixel, box 320x320 pixel
180°
3.47 Å
G
Import
10,053 micrographs
Blob Picker
(1,9863,191)
Extract
2.090 Å/pixel
box 250x250 pixel
(1,431,545)
2D Classification
snoRNP (60,780)
Topaz 8x binned
(902,527)
2.090 Å/pixel
box 160x160 pixel
Heterogeneous Refinement
(289,037)
(290,114)
(323,376)
(70,449)
(182,190)
(70,737)
Topaz 8x binned
(1,789,289)
150 Å
2.090 Å/pixel
box 160x160 pixel
Heterogeneous Refinement
(619,171)
(551,375)
(618,743)
Non-uniform Refinement
Extract
1.045 Å/pixel
box 320x320 pixel
(618,214)
3D Classification
Non-uniform Refinement
(314,442)
(303,772)
Heterogeneous Refinement
Heterogeneous Refinement
(183,000)
(63,805)
(67,637)
(71,225)
(66,095)
(166,452)
(6,478)
(5,298)
(171,224)
(126,953)
(20,504)
(18,995)
Local Filtering
180°
Non-uniform Refinement
2.89 Å
snR30 snoRNP
Utp23-Krr1ΔC3
State1
Local Refinement
2.99 Å
3D Classification
Non-uniform Refinement
(59,660)
(67,293)
Local Filtering
3.18 Å
snR30 snoRNP
Utp23-Krr1ΔC3
State2
H
2D Classes
snR30 snoRNP
Utp23-Krr1ΔC3 State1
150 Å

**Figure S4 (related to Figure 3 and Figure 6). Cryo-EM sorting schemes of the Utp23-Krr1 wt and ΔC3 datasets.** (**A**, **B** and **D**, **E**) Representative electron micrographs (**A**, **D**) and initial 2D class averages (**B**, **E**) of the Utp23-Krr1 wt (**A**, **B**) and the Utp23-Krr1ΔC3 (**D**, **E**) purifications. (**C**, **F**, **G**) Processing schemes of the Utp23-Krr1 wt (**C**) and Utp23-Krr1ΔC3 (**F**, **G**) datasets. The Krr1 wt dataset was exclusively processed with CryoSPARC, while for the Utp23-Krr1ΔC3 Relion and CryoSPARC were used. (**G**) Cryo-EM processing and sorting scheme of the Utp23-Krr1ΔC3 dataset used to obtain the final cryo-EM density maps of snR30 snoRNP Utp23-Krr1ΔC3 State1 and State2. Processing was performed with CryoSPARC. (**C**, **F**, **G**) The final maps are highlighted, particle numbers are shown in brackets and the applied pixel sizes and box sizes are indicated. For details see Material and Methods section. (**H**) Main 2D class averages for the final snR30 snoRNP Utp23-Krr1ΔC3 State1.

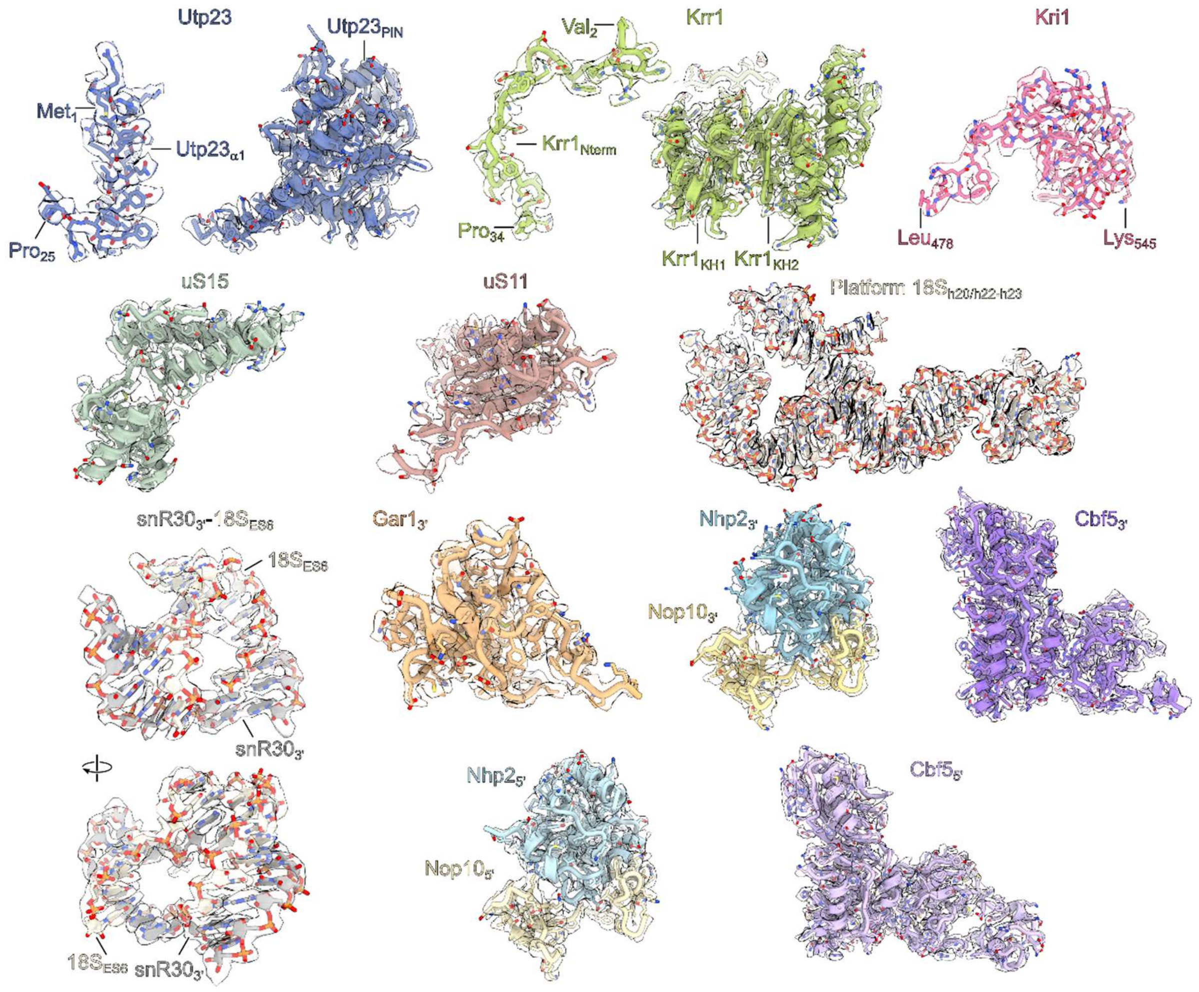


**Figure S5 (related to Figure 3). Cryo-EM densities and models of snR30 snoRNP factors.** Molecular models and corresponding segmented cryo-EM density maps of the factors identified within the snR30 snoRNP Utp23-Krr1∆C3 State 1.

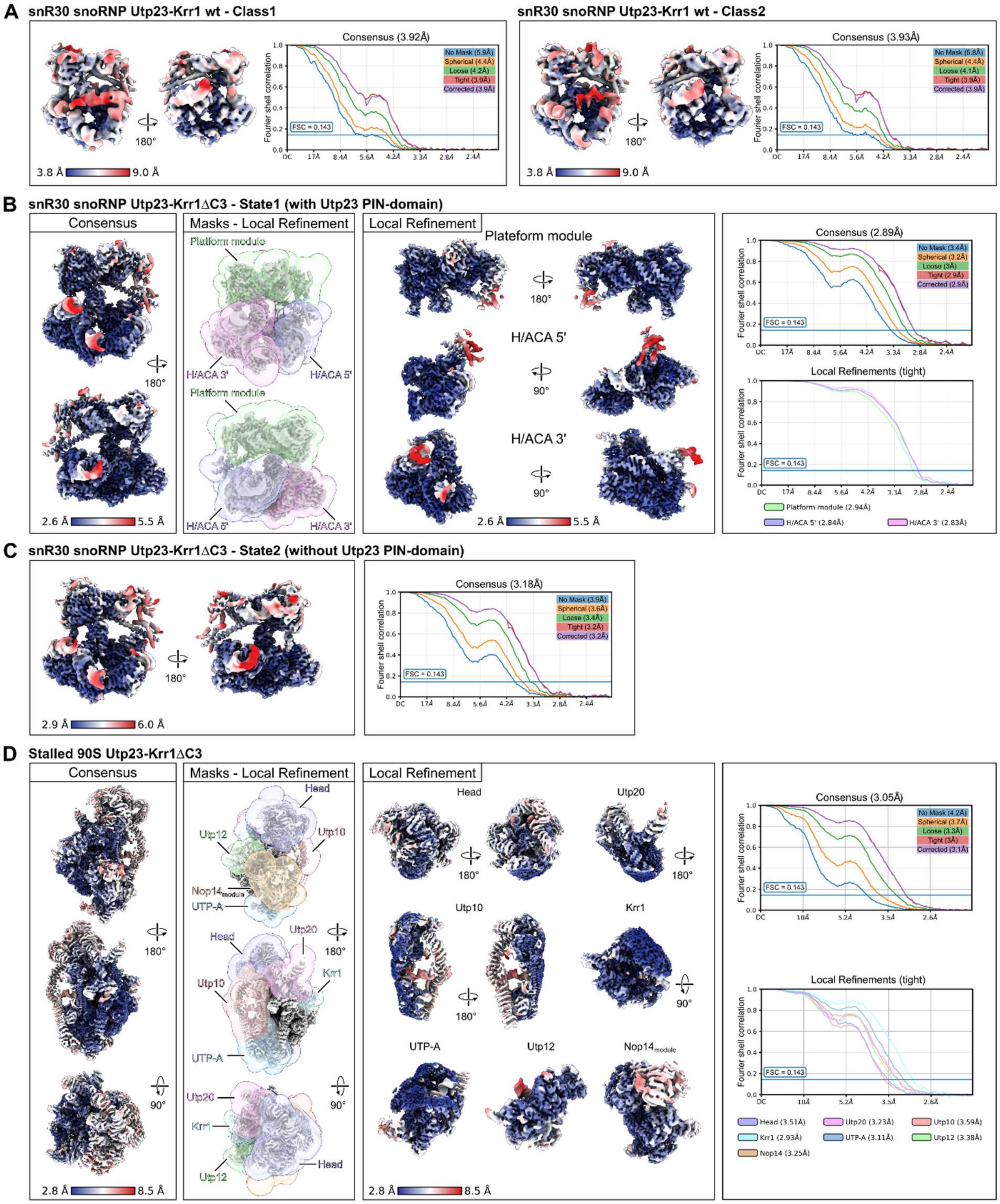


**Figure S6 (related to Figure 3 and Figure 6). Local Resolution and Fourier shell correlation curves.** (**A**) Local resolution filtered cryo-EM density maps and Fourier shell correlation (FSC) curves of the snoRNP classes obtained from the Utp23-Krr1 wt dataset. (**B**, **C**) Local filtered cryo-EM density maps colored according to local resolution and FSC curves of the snoRNP states of the Utp23-Krr1ΔC3 dataset.

For State 1 (**B**) masks applied for local refinements and the corresponding local refined maps colored according to the local resolution are shown (middle panels). (**D**) Local resolution filtered consensus map of the stalled 90S particle (Utp23-Krr1∆C3 dataset, left panel). Masks for local refinements and cryo-EM density maps of the local refinements colored according to local resolution (middle). FSC curves of the consensus refinement (upper right) and FSC curves of the individual local refinements (lower right).

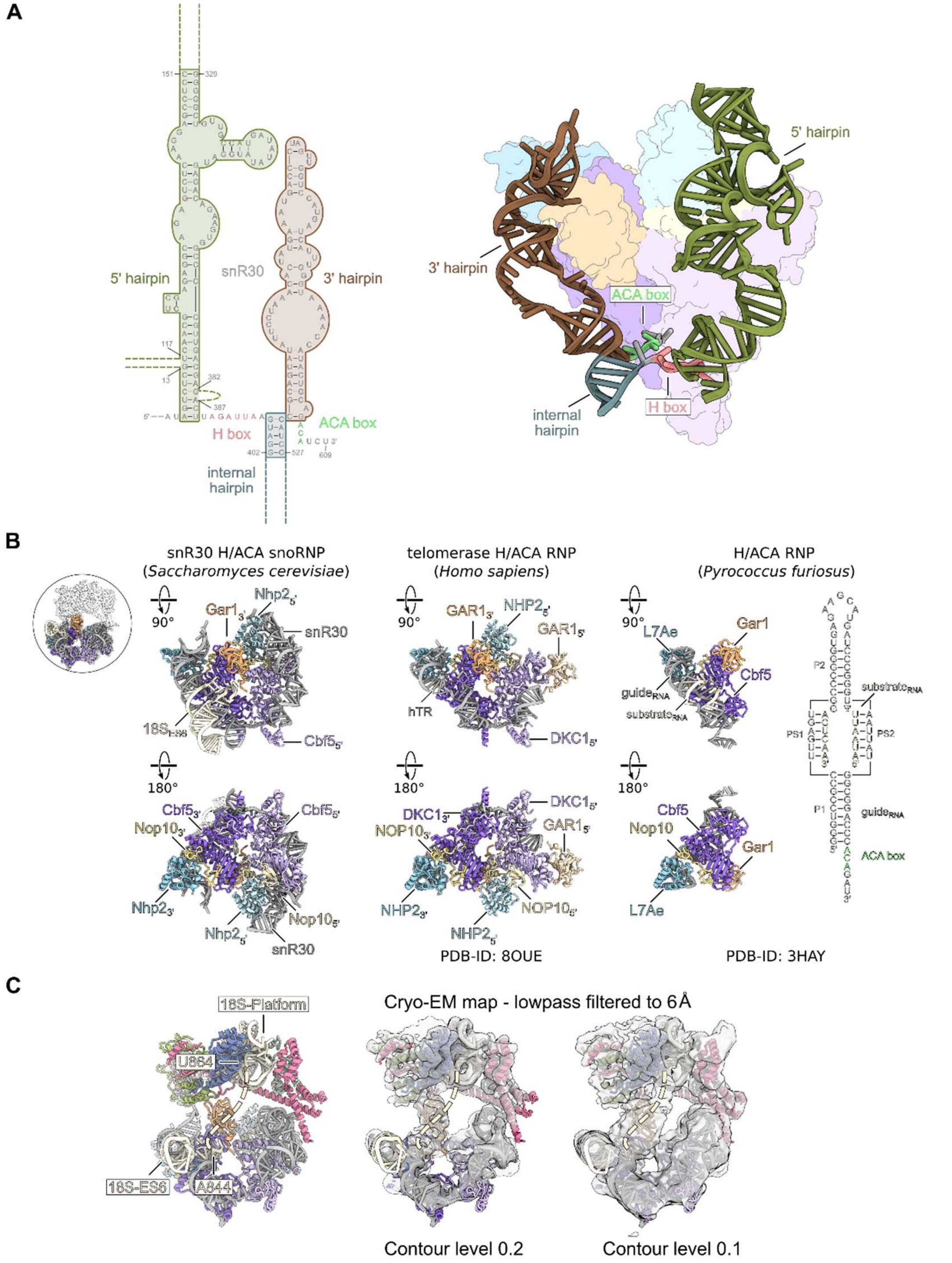


**Figure S7 (related to Figure 4). snR30 structure and comparison of the H/ACA module.** (**A**), Secondary structure of the modelled snR30 areas. The 5', 3' and internal hairpins as well as the H and ACA box consensus sequences are highlighted (left panel). Molecular model of the snR30 snoRNA and surface views of the H/ACA core proteins (right panel). (**B**) Comparison of the snR30 H/ACA module with the human telomerase H/ACA module (PDB-ID: 8OUE) and the archaea H/ACA RNP from *Pyrococcus furiosus* (PDB-ID: 3HAY). The secondary structures of the archaea guide and substrate RNAs are shown. (**C**) Model of the snR30 snoRNP and the corresponding lowpass filtered cryo-EM map at the indicated contour levels. The connecting density between the 18S-ES6 bound to the snR30 3' hairpin and 18S-Platform rRNA h22 is visible at lower contour levels and indicated as dashed line. The boundaries of the 18S molecular model are indicated (left).

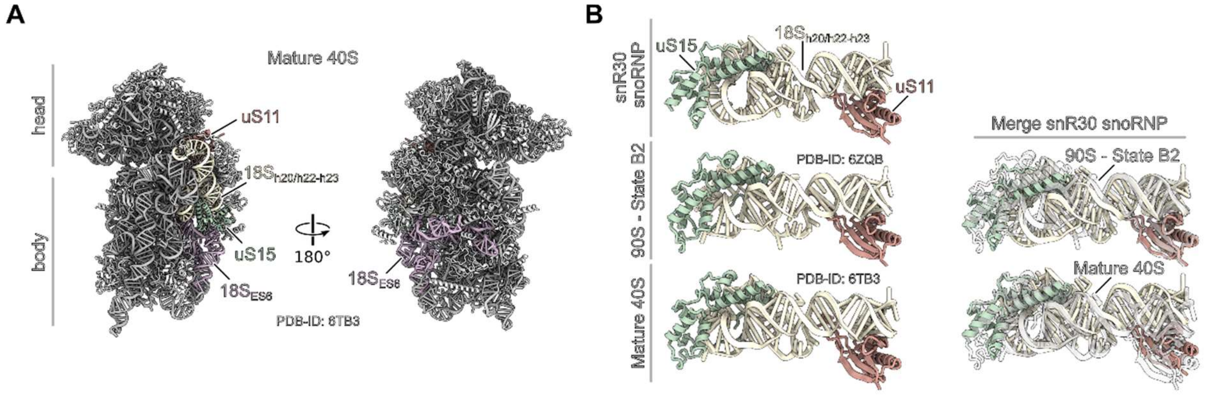


**Figure S8 (related to Figure 5). The platform module has a close to mature conformation.** (**A**) Model of the mature 40S subunit (PDB-ID: 6TB3). Platform helices h20, h22-h23, the complete ES6 and ribosomal proteins uS11 and uS15 are colored and labeled. (**B**) Comparison of the platform module within the snR30 snoRNP (Utp23-Krr1ΔC3 State 1), when bound to the 90S particle (State B2, PDB-ID: 6ZQB) and within the mature 40S subunits (PDB-ID:6TB3) (left panels). Molecular models were rigid body fitted. The snR30 snoRNP platform model is colored and 90S or 40S models are shown in transparent gray (right panels).

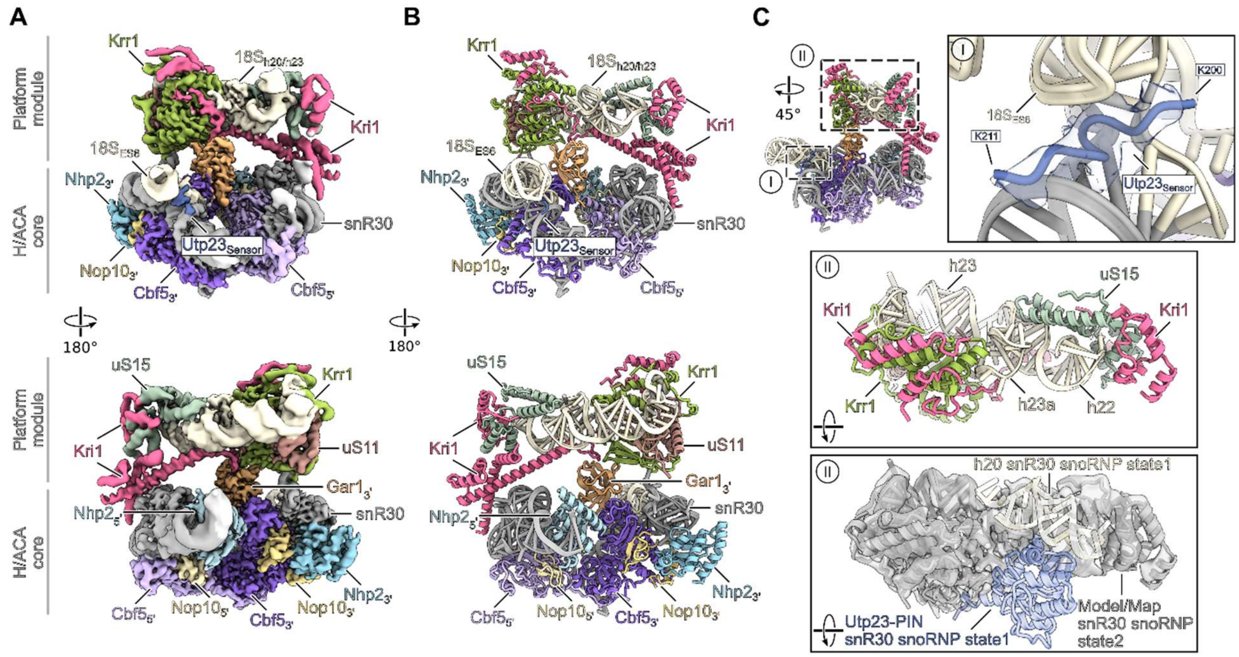


**Figure S9 (related to Figure 6). snR30 snoRNP lacking the bound Utp23 PIN domain.** (**A**, **B**) Colored cryo-EM map (**A**) and molecular model (**B**) of the snR30 snoRNP Utp23-Krr1ΔC3 State 2 without a stable bound Utp23-PIN domain. Bound factors, the H/ACA core and the platform module are labeled. (**C**) Overview and close-ups of State 2 focusing on the Utp23 interaction regions (upper left). (I) Model of the snR30:18S-ES6 three-way junction and cryo-EM density and model of the Utp23-Sensor (aa200-211) are shown. (II) Top view showing the molecular model of the snR30 snoRNP Utp23-Krr1ΔC3 State 2 platform module (upper panel) and overlay with the models of 18S rRNA h20 and Utp23-PIN domain of snR30 snoRNP Utp23-Krr1ΔC3 State 1. Model and density of State 2 are shown in gray and the h20 and the Utp23-PIN domain of State 1 are displayed as transparent colored molecular model (lower panel).

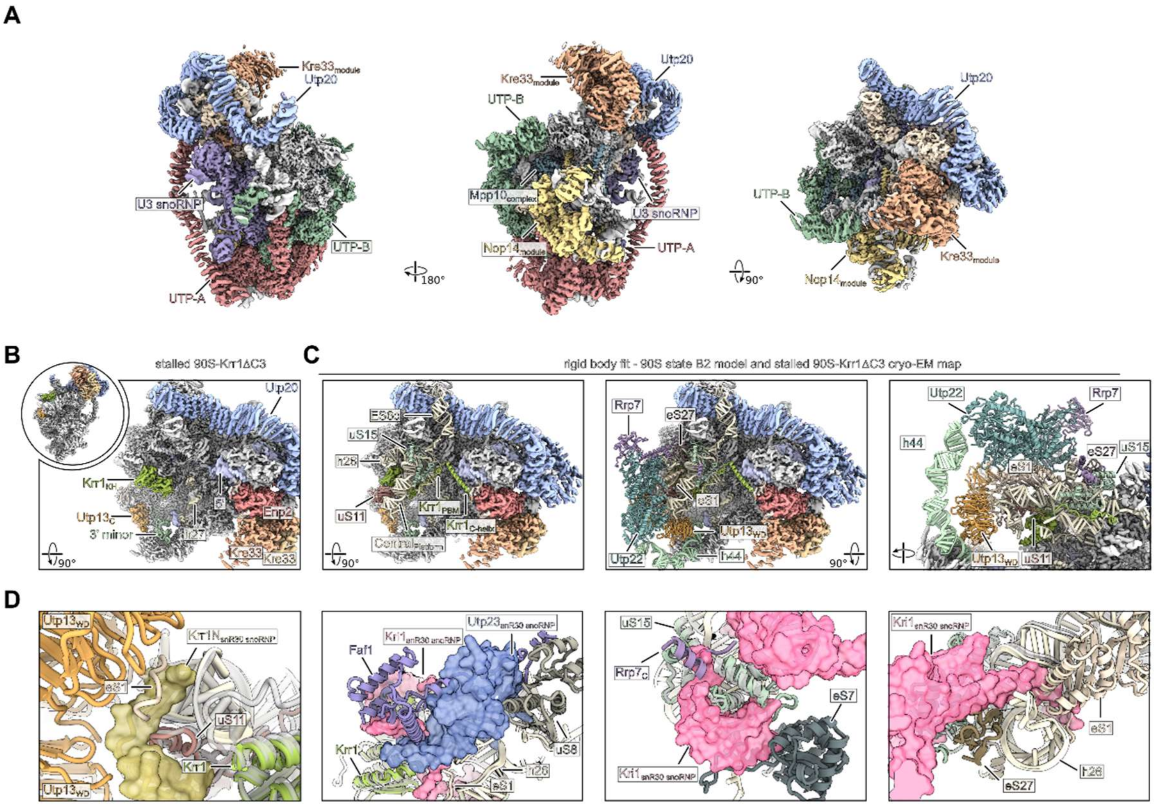


**Figure S10 (related to Figure 6). Structure of the stalled 90S and comparison with the native 90S state B2.** (**A**) Cryo-EM density map of the stalled 90S state purified through Utp23-Krr1ΔC3. AFs and 90S biogenesis modules are colored and labeled. (**B**) Overview of the stalled 90S and close-up of the immature central domain region. Depicted factors are labeled for orientation. A Krr1 molecule was identified in its previously observed position within the stalled 90S particle which could be a wild-type copy of Krr1 (note that in the purification of the dominant-negative *krr1*ΔC3 mutant, wild-type *KRR1* is still present). (**C**) Overlay with the molecular model of the state B2 90S (PDB-ID: 6ZQB) showing the central domain rRNA and ribosomal proteins uS11 and uS15 (left panel) and in addition the interacting UTP-C module (Utp22-Rrp7), the Utp13 WD propellers and h44 of the 18S 3' minor domain (middle and right panel). (**D**) Magnification views highlighting the steric clashes between the snR30 snoRNP platform module and the 90S. The snR30 snoRNP and the 90S state B2 (PDB-ID: 6ZQB) models were rigid body fitted. Molecular models of the snR30 snoRNP are shown as transparent surface and transparent gray models. The 90S model is depicted as molecular model and factors clashing with the snR30 snoRNP structure are coloured and labeled.
