## Supplementary material for "H/ACA snoRNP guides ribosomal RNA subdomain folding in a satellite particle before joining the core 90S pre-ribosome": Key Resources Table

| REAGENT or RESOURCE | SOURCE | IDENTIFIER |
|---|---|---|
| **Antibodies** | | |
| Anti-FLAG | SIGMA-Aldrich | A8592;<br>RRID: AB_439702 |
| Anti-ProteinA | SIGMA-Aldrich | P1291; RRID: AB_260996 |
| **Bacterial and virus strains** | | |
| *Escherichia coli* DH5α | | |
| **Chemicals, peptides, and recombinant proteins** | | |
| FLAG Peptide | CASLO | N/A |
| TEV protease | Parks et al 1994 | N/A |
| SYBR Green II RNA gel stain | SIGMA-Aldrich | S9305 |
| SIGMA*FAST* | SIGMA-Aldrich | S8830 |
| RiboLock RNase inhibitor | Thermo Scientific | EO0381 |
| T4 PNK | NEB | M0201 |
| **Critical commercial assays** | | |
| ANTI-FlagM2 Affinity Gel | SIGMA-Aldrich | A2220 |
| IgG–Sepharose 6 Fast Flow | GE Healthcare | 17096902 |
| **Deposited data** | | |
| Raw and analyzed data | This study | xxxMENDELEYxxx |
| RAW files of mass spectrometry dataset 1 (Figure 1e) | This study | ProteomeXchange: 10.6019/PXD054031 |
| RAW files of mass spectrometry dataset 2 (Figure 2b) | This study | ProteomeXchange: 10.6019/PXD054032 |
| RAW files of mass spectrometry dataset 3 (Figure 2d) | This study | ProteomeXchange: 10.6019/PXD054034 |
| RAW files of mass spectrometry dataset 4 (Figure 3c) | This study | ProteomeXchange: 10.6019/PXD054035 |

| | | |
|---|---|---|
| snR30 snoRNP Utp23 Krr1ΔC3 State1 | This study | DB-9G25, EMD-50964, EMD-50958, EMD-50959, EMD-50960. EMD-50961 |
| snR30 snoRNP Utp23-Krr1ΔC3 State2 | This study | EMD-50968, PDB-9G28 |
| Stalled 90S Utp23-Krr1ΔC3 | This study | PDB-9G33, EMD-50991, EMD-50647, EMD-50648, EMD-50649, EMD-50650, EMD-50651, EMD-50652, EMD-50653, EMD-50654 |
| snR30 snoRNP Utp23-Krr1 wt Class1 | This study | EMD-50967 |
| snR30 snoRNP Utp23-Krr1 wt Class2 | This study | EMD-50969 |
| **Experimental models: Organisms/strains** | | |
| *ade2-1 ,trp1-1, leu2-3,112, his3-11,15, ura3-1, can1-100* | [1] | W303 |
| W303, *krr1*::*HIS3*, [pRS316 *KRR1*] | [2] | Krr1 shuffle |
| W303, *krr1*::*HIS3*, *utp7*::*hphNT1* [pRS316 *KRR1*, pRS316 *UTP7*] | This study | Krr1-Utp7 double shuffle |
| W303, *krr1*::*HIS3*, *utp7*::*hphNT1*, *NOC4*-FLAG-TEV-ProtA::*natNT2* [pRS316 *KRR1*, pRS316 *UTP7*] | This study | Krr1-Utp7 double shuffle, Noc4-FTpA |
| W303, *krr1*::*HIS3*, *UTP10*-FLAG-TEV-ProtA::*kanMX6* [pRS316 *KRR1*] | This study | Krr1 shuffle, Utp10-FTpA |
| W303, *krr1*::*HIS3*, *UTP7*-FLAG-TEV-ProtA::*natNT2* [pRS316 *KRR1*] | This study | Krr1 shuffle, Utp7-FTpA |
| **Oligonucleotides** | | |
| 5′-TTATGGGACTTGTT-3′ | [3] | anti-U3 |
| 5′-TCACTCAGACATCCTAGG-3′ | [4] | anti-U14 |
| 5′-ATGTCTGCAGTATGGTTTTAC-3′ | [5] | anti-snR30 |
| **Recombinant DNA** | | |
| $P_{KRR1}$-*KRR1*-$T_{ADH1}$, *URA3*, ARS/CEN, *AmpR* | [2] | pRS316 Krr1wt |
| $P_{KRR1}$-*KRR1*-$T_{ADH1}$, *TRP1*, ARS/CEN, *AmpR* | [2] | pRS314 Krr1wt |
| $P_{KRR1}$-*krr1* ΔC-$T_{ADH1}$, *TRP1*, ARS/CEN, *AmpR* | [2] | pRS314 Krr1ΔC |
| $P_{KRR1}$-*krr1* ΔC1-$T_{ADH1}$, *TRP1*, ARS/CEN, *AmpR* | This study | pRS314 Krr1ΔC1 |

| | | |
|---|---|---|
| $P_{KRR1}$-*krr1* ΔC2-$T_{ADH1}$, *TRP1*, ARS/CEN, *AmpR* | This study | pRS314 Krr1ΔC2 |
| $P_{KRR1}$-*krr1* ΔC3-$T_{ADH1}$, *TRP1*, ARS/CEN, *AmpR* | This study | pRS314 Krr1ΔC3 |
| $P_{KRR1}$-*krr1* ΔN-$T_{ADH1}$, *TRP1*, ARS/CEN, *AmpR* | This study | pRS314 Krr1ΔN |
| $P_{UTP7}$-*UTP7*-$T_{ADH1}$, *URA3*, ARS/CEN, *AmpR* | This study | pRS316 Utp7 |
| $P_{UTP7}$-*UTP7*-$T_{ADH1}$, *LEU2*, ARS/CEN, *AmpR* | This study | pRS315 Utp7 |
| $P_{UTP7}$-*UTP7*-FLAG-TEV-ProtA-$T_{ADH1}$, *LEU2*, ARS/CEN, *AmpR* | This study | pRS315 Utp7-FTpA |
| $P_{GAL1-10}$-MCS-$T_{ADH1}$, *LEU2*, 2μ, *AmpR* | This study | pGAL-empty |
| $P_{GAL1-10}$-*KRR1*-$T_{ADH1}$, *LEU2*, 2μ, *AmpR* | This study | pGAL-Krr1 Leu |
| $P_{GAL1-10}$-*KRR1*-$T_{ADH1}$, *TRP1*, 2μ, *AmpR* | This study | pGAL-Krr1 Trp |
| $P_{GAL1-10}$-*krr1* ΔC3-$T_{ADH1}$, *LEU2*, 2μ, *AmpR* | This study | pGAL-Krr1ΔC3 Leu |
| $P_{GAL1-10}$-*krr1* ΔC3-$T_{ADH1}$, *TRP1*, 2μ, *AmpR* | This study | pGAL-Krr1ΔC3 Trp |
| $P_{GAL1-10}$-*krr1*-$T_{ADH1}$, *TRP1*, 2μ, *AmpR* | This study | pGAL-Krr1ΔN Trp |
| **Software and algorithms** | | |
| ClustalOMEGA | EMBL-EBI | https://www.ebi.ac.uk/Tools/msa/clustalo |
| USCF ChimeraX | [6] | http://www.cgl.ucsf.edu/chimerax |
| Coot | [7] | https://www2.mrc-lmb.cam.ac.uk/personal/pemsley/coot/ |
| PSIPRED | | http://bioinf.cs.ucl.ac.uk/psipred/ |
| MaxQUANT | [8] | https://www.maxquant.org |
| EM-TOOLS | TVIPS | https://www.tvips.com/imaging-software/em-tools/ |
| MotionCor2 | [9] | https://emcore.ucsf.edu/cryoem-software |
| CTFFIND4 | [10] | http://grigoriefflab.janelia.org/ctffind4 |
| CryoSPARC | [11] | |
| Relion 3.1.3 | [12] | |
| PHENIX | [13] | https://www.phenix-online.org |
| Alphafold | [14] | Alphafold |

| | |
|---|---|
| Alphafold multimer | doi: https://doi.org/10.1101/2021.10.04.463034 |
| **Other** | |
| Grids | Quantifoil Micro Tools GmbH |

1 Thomas, B. J. & Rothstein, R. Elevated recombination rates in transcriptionally active DNA. *Cell* **56**, 619-630 (1989). https://doi.org:10.1016/0092-8674(89)90584-9

2 Cheng, J. *et al.* Thermophile 90S Pre-ribosome Structures Reveal the Reverse Order of Co-transcriptional 18S rRNA Subdomain Integration. *Mol Cell* **75**, 1256-1269 e1257 (2019). https://doi.org:10.1016/j.molcel.2019.06.032

3 Sharma, K. & Tollervey, D. Base pairing between U3 small nucleolar RNA and the 5' end of 18S rRNA is required for pre-rRNA processing. *Mol Cell Biol* **19**, 6012-6019 (1999). https://doi.org:10.1128/MCB.19.9.6012

4 Allmang, C. *et al.* Functions of the exosome in rRNA, snoRNA and snRNA synthesis. *EMBO J* **18**, 5399-5410 (1999). https://doi.org:10.1093/emboj/18.19.5399

5 Fath, S. *et al.* Association of yeast RNA polymerase I with a nucleolar substructure active in rRNA synthesis and processing. *J Cell Biol* **149**, 575-590 (2000). https://doi.org:10.1083/jcb.149.3.575

6 Pettersen, E. F. *et al.* UCSF ChimeraX: Structure visualization for researchers, educators, and developers. *Protein Sci* **30**, 70-82 (2021). https://doi.org:10.1002/pro.3943

7 Emsley, P., Lohkamp, B., Scott, W. G. & Cowtan, K. Features and development of Coot. *Acta Crystallogr D Biol Crystallogr* **66**, 486-501 (2010). https://doi.org:10.1107/S0907444910007493

8 Cox, J. & Mann, M. MaxQuant enables high peptide identification rates, individualized p.p.b.-range mass accuracies and proteome-wide protein quantification. *Nat Biotechnol* **26**, 1367-1372 (2008). https://doi.org:10.1038/nbt.1511

9 Zheng, S. Q. *et al.* MotionCor2: anisotropic correction of beam-induced motion for improved cryo-electron microscopy. *Nat Methods* **14**, 331-332 (2017). https://doi.org:10.1038/nmeth.4193

10 Rohou, A. & Grigorieff, N. CTFFIND4: Fast and accurate defocus estimation from electron micrographs. *J Struct Biol* **192**, 216-221 (2015). https://doi.org:10.1016/j.jsb.2015.08.008

11 Punjani, A., Rubinstein, J. L., Fleet, D. J. & Brubaker, M. A. cryoSPARC: algorithms for rapid unsupervised cryo-EM structure determination. *Nat Methods* **14**, 290-296 (2017). https://doi.org:10.1038/nmeth.4169

12 Zivanov, J. *et al.* New tools for automated high-resolution cryo-EM structure determination in RELION-3. *Elife* **7** (2018). https://doi.org:10.7554/eLife.42166

13 Liebschner, D. *et al.* Macromolecular structure determination using X-rays, neutrons and electrons: recent developments in Phenix. *Acta Crystallogr D Struct Biol* **75**, 861-877 (2019). https://doi.org:10.1107/S2059798319011471

14 Jumper, J. *et al.* Highly accurate protein structure prediction with AlphaFold. *Nature* **596**, 583-589 (2021). https://doi.org:10.1038/s41586-021-03819-2
